# Cortical surface alterations in cingulate and frontal regulatory areas underlie insensitive mothering

**DOI:** 10.1101/2020.01.29.924613

**Authors:** Inmaculada León, María José Rodrigo, Ileana Quiñones, Juan Andrés Hernández-Cabrera, Lorna García-Pentón

## Abstract

This study focuses on severe insensitive or neglectful mothering, the most prevalent type of child maltreatment, to examine cortical surface feature alterations underlying maternal functioning and their impact on mother-child interactive bonding. High-resolution 3D volumetric images were obtained on 24 neglectful (NM) and 21 non-neglectful control (CM) mothers. Using surface-based morphometry, we compared differences in cortical thickness and surface area. Mothers completed alexithymia and cortical integrity measures and participated with their children in a play task (Emotional Availability Scale). We found cortical thinning for NM in the right rostral middle frontal gyrus and the right anterior/medial cingulate cortex, and also increased surface area in the right occipital lingual and fusiform areas and the caudal middle frontal area. Mediation analyses showed that cognitive integrity and alexithymia mediated, respectively, the positive and negative effect of the rostral middle frontal gyrus on Emotional Availability. The findings suggest cortical thinning in the rostral frontal area underlying high-order regulatory functioning as being critical for poor maternal self-awareness of emotions and the organization of coordinated actions during mother-child interactive bonding.

## INTRODUCTION

The sensitive caregiving response in mother-child interactive bonding is a critical aspect of adequate childrearing, since it is related to the child’s secure attachment and healthy development (Weinfield et al., 2008). The quality of this maternal response is the first factor contributing to the development of the neural mechanisms of empathy in children (Levy, Goldstein, & Feldman, 2019). Although most mothers can respond appropriately to their child’s needs, extreme cases of insensitive caregiving do exist. Maternal neglect, the most common form of child maltreatment, involves a drastic failure to provide the child with food, clothing, shelter, medical care, supervision or emotional support that places the child’s safety at risk (Petersen et al., 2014; Stoltenborgh et al., 2013) and entails negative behavioral, neurobiological and clinical consequences for the offspring (Nemeroff, 2016; Teicher et al., 2016). Despite its importance, we still know little about the neuroanatomical bases underlying neglectful (vs. sensitive) caregiving, limiting our understanding of the associations between brain structure and human mothering.

It is well known that the maternal brain exhibits higher activations to the mother’s own versus an unfamiliar child’s signals in empathy-related regions (such as the anterior insula, supplementary motor area, cingulate cortex and inferior frontal regions), which is crucial for sensitive responses to the child (see reviews in (Feldman, 2015; Kim et al., 2016; H. Rutherford & Mayes, 2016; Swain et al., 2007). Conversely, the neglectful mother’s brain shows reduced activations to infant crying faces in frontal and anterior cingulate areas (León et al., 2019). We recently observed in a voxel-based morphometry (VBM) study that grey matter (GM) volume decreased in the right insula, anterior/middle cingulate and inferior/middle frontal gyrus and increased in the right fusiform and cerebellum for neglectful as compared to non-neglectful mothers (Rodrigo et al., 2019). However, GM volume is an unspecific measure that can be influenced by different combinations of independent anatomical factors, such as cortical surface area (SA) and cortical thickness (CT) (Mechelli et al., 2005; Rimol et al., 2012; Winkler et al., 2010). SA and CT are morphological features of the cerebral cortex that follow different genetic, phylogenetic and developmental routes, capturing better the anatomical and functional variability in the brain (Winkler et al., 2012). They offer more sensitive and more specific quantitative indices to predict brain alterations (Winkler et al., 2017). Specifically, CT is influenced by the number and size of the cells, organized vertically within a cortical column (Buxhoeveden & Casanova, 2002), while SA is driven by the number of columns, organized horizontally across the cortical surface, as well as by the number of connections and the extent of their myelination (Goldman-Rakic, 1995; Lyttelton et al., 2009). In the current study, we take advantage of a surface-based approach over the VBM (Van Essen et al., 1998) to further refine the analysis of the cortical morphology in sensitive caregiving response.

Our main goal is to identify the specific cortical morphometric patterns that are altered in neglectful mothering, as a way to better understand the underlying mechanism associated with poor mother-child bonding interactions. This goal involved two specific objectives: i) to identify cortical thickness and surface area differences between a group of 24 neglectful mothers (NM) and a control group of 21 non-neglectful mothers (CM) who are socio-demographically similar, and ii) to determine the potential role of the altered cortical features by ascertaining their correlation with behavioral measurements relevant for effective mother-child interactive bonding (brain structure-behavior associations). Greater cortical thickness in the prefrontal cortex, the lateral occipital areas and the fusiform gyri has been associated with later postpartum months in first-time mothers, with the cortical increase in the prefrontal cortex uniquely associated with higher levels of parental self-efficacy (Kim et al., 2018). Based on these results as well as on previous evidence from the VBM study with NM (Rodrigo et al., 2019), we expected to find CT and SA group differences mainly in frontal, cingulate and occipital areas.

We examined the brain structure-behavior association in NM using Emotional Availability (EA), measured through a mother-child play task, as a proxy for the quality of mother-child bonding interactions (Biringen, 2000; Biringen et al., 2014). Quality of EA is a crucial feature of the dyadic functioning that is predictive of the mother’s reported child attachment (Altenhofen et al., 2013). Performing the play task requires sensitive and responsive dyadic exchanges (i.e., sensitivity score that involves parents having a calm emotional presence and reading a child’s emotional cues appropriately). The play task also requires the organization of coordinated actions towards achieving joint goals (i.e., structuring score that involves parental guidance, mentoring and empowerment of the child’s autonomous pursuits). In this regard, synchrony is a crucial feature in mother-child interactive bonding that is related to attachment quality (Beebe & Steele, 2013; Biro et al., 2017). Accordingly, NM have been found to exhibit lower capacities to perform the interactive play task (Rodrigo et al., 2019).

Some differences in parental traits, like alexithymia, which affects their ability to be aware of their own emotional state to empathically respond to the feelings of the child (Decety & Jackson, 2004), or cognitive capacity, which allows them to attend, plan, memorize and execute actions (Friedman et al., 2008; Schmeichel & Tang, 2014), may be involved in the effective performance of the play task. Alexithymia, on the one hand, is a personality trait associated with metacognitive impairments in emotional processing that involves a difficulty in recognizing and describing one’s emotions, differentiating mental states from bodily sensations, and minimizing emotional experience by focusing attention externally (Herbert et al., 2011; Mantani et al., 2005; Taylor, 2000). Cognitive integrity, on the other hand, is a global construct of effective cognitive performance that mainly comprises attention, calculus, memory and language processes (Folstein, Folstein, McHugh et al., 1975). Previous results of our group have shown that NM scored lower than CM in alexithymia and cognitive integrity measures (Herrero-Roldán et al., 2019; Rodrigo et al., 2016).

Accordingly, we examined whether alexithymia and cognitive integrity may play a mediating role in the brain structure-EA associations, since they are traits related to brain areas that may be affected in NM and could be involved in the effective performance of the mother-child play task. Specifically, alexithymia is associated with reduced neural responses in the amygdala, insula and anterior/posterior cingulate cortex, as well as in the dorsomedial prefrontal cortex (Moriguchi & Komaki, 2013; van der Velde et al., 2013). Moreover, high alexithymic compared to low alexithymic women showed reduced activations in the orbito-frontal cortex (OFC) and anterior cingulate cortex (ACC) during empathizing with an imagined 1-year-old child (Lenzi et al., 2013). NM have also shown a poor ability to express their feelings verbally (Herrero-Roldán et al., 2019; Polansky, 1985). Therefore, we expected that introducing alexithymia as a mediator would help enhance associations between the cortical alterations observed and EA.

Cognitive integrity is used in prevention and clinical practice to help distinguish normal variability in healthy adults from several degrees of pathological deviations (Mitchell, 2009). Higher cognitive integrity has been positively correlated with the GM volume bilaterally in frontal regions (Arlt et al., 2013), as well as in the limbic and cingulate regions (Dinomais et al., 2016). Studies showed that mothers with poorer working memory, a main component of cognitive integrity, had worse cognitive control of their emotions, thoughts and behaviors during a frustrating cooperation task with their child and were more likely to show reactive negativity in response to children’s challenging behaviors (Deater-Deckard et al., 2010). Since NM have shown poor cognitive integrity (Rodrigo et al., 2016), we also expected that cognitive integrity as a mediator variable would enhance brain structure-EA associations.

Overall, this is the first study looking for cortical features underlying neglectful mothering and their relationships with emotional and cognitive parental traits that could be relevant for mother-child interactive bonding.

## Methods

### Participants

Forty-five mothers (24 NM and 21 CM) voluntarily participated in the experiment. They all were recruited through the same Primary Health Center in Tenerife, Spain. Written informed consent was obtained from all the participants and the Ethics Committee of the University of La Laguna approved the study’s protocol. General inclusion criteria were being the biological mother of a child under three years old who had not been placed in foster care at any point in their history nor been born prematurely or suffered perinatal or postnatal medical complications, according to the pediatricians’ reports. Specific inclusion criteria for the neglectful mother group were a substantiated case of neglect registered in the last 12 months by Child Protective Services (CPS) and complying with the indicators of the Maltreatment Classification System (MCS) for severe neglect (Barnett et al., 1993). Inclusion criteria for the control group were negative scores in all the MCS neglect indicators and the absence of CPS or Preventive Services records for the family. As for the sociodemographic profile of mothers, they were all in their early 30s, had a similar mean age of the target child, lived in rural areas and shared a similar low level of education; the NM were more likely than the CM to live in single-parent families and to receive financial assistance (Table 1). According to the neglect risk profile rated by the social workers, most mothers in the neglectful group had a history of childhood maltreatment or neglect and scored positively in neglectful caregiving indicators [see Supplementary *Methods* 1 for participant recruitment and procedure, and 2 for risk profile measures].

**Table 1.**
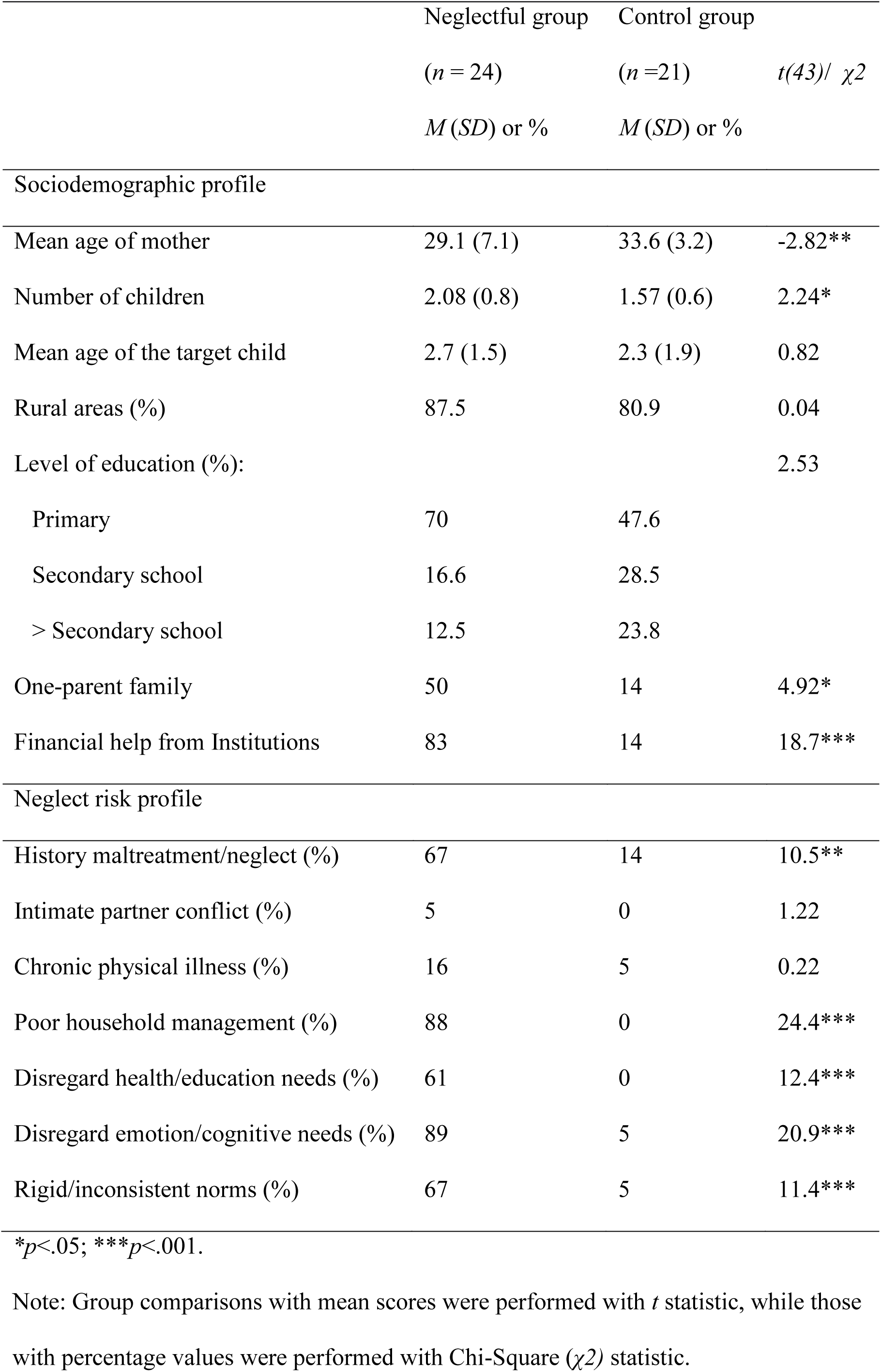
Sociodemographic and neglect risk profile in Neglectful and Control Groups

| | Neglectful group<br>(*n* = 24)<br>*M* (*SD*) or % | Control group<br>(*n* =21)<br>*M* (*SD*) or % | *t*(43)/ $\chi^2$ |
| --- | --- | --- | --- |
| Sociodemographic profile |  |  |  |
| Mean age of mother | 29.1 (7.1) | 33.6 (3.2) | -2.82** |
| Number of children | 2.08 (0.8) | 1.57 (0.6) | 2.24* |
| Mean age of the target child | 2.7 (1.5) | 2.3 (1.9) | 0.82 |
| Rural areas (%) | 87.5 | 80.9 | 0.04 |
| Level of education (%): |  |  | 2.53 |
| Primary | 70 | 47.6 |  |
| Secondary school | 16.6 | 28.5 |  |
| > Secondary school | 12.5 | 23.8 |  |
| One-parent family | 50 | 14 | 4.92* |
| Financial help from Institutions | 83 | 14 | 18.7*** |
| Neglect risk profile |  |  |  |
| History maltreatment/neglect (%) | 67 | 14 | 10.5** |
| Intimate partner conflict (%) | 5 | 0 | 1.22 |
| Chronic physical illness (%) | 16 | 5 | 0.22 |
| Poor household management (%) | 88 | 0 | 24.4*** |
| Disregard health/education needs (%) | 61 | 0 | 12.4*** |
| Disregard emotion/cognitive needs (%) | 89 | 5 | 20.9*** |
| Rigid/inconsistent norms (%) | 67 | 5 | 11.4*** |

\**p*<.05; \*\*\**p*<.001.
Note: Group comparisons with mean scores were performed with *t* statistic, while those with percentage values were performed with Chi-Square ( $\chi^2$ ) statistic.

### Behavioral and personality measures

*Mini-mental state examination* (MMS; Folstein, Folstein, & McHugh, 1975); the Spanish version (Blesa et al., 2001) of the MMS was administered to assess differences in cognitive function. In 11 items, the MMS records temporal and spatial orientation; attention, concentration and memory capacity; abstraction capacity (calculation); language ability and visuospatial perception; and ability to follow basic instructions, giving a final accumulative score. CI was lower in NM than in CM, respectively (*M* = 25.9; *SD* = 1.92; *M* = 28.7; *SD* = 1.42; *t*(43) = 5.4; *p* = .000; *δ* = 1.61).

The *Mini International Neuropsychiatric Interview* (M.I.N.I. 6.0; Spanish version Ferrando et al., 2000), including 15 major psychiatric disorders, was administered [see Supplementary *Methods* 3, Table S1]. The two groups mainly differed in five psychopathological variables, marked in italics, which survived the Bonferroni correction and were submitted to a Principal Component Analysis. Results gave one factor solution: “Psychiatric Disorders” (PD), with moderate inter-correlations among the five variables, KMO = 0.65, Eigenvalue = 2.53, with an explained variance of 51%, with the coefficient scores in PD being higher in NM (*M* = 0.67, *SD* = 0.99) than in CM (*M* = −0.69, *SD* = 0.23), *t*(43) = 6.35; *p* = .000; *δ* = 1.90. None of the mothers in either group was being medicated for psychiatric disorders at the time of testing. The coefficient score in PD was used as a regressor in the SPM model to control as much as possible for its effect on brain volume differences. Given that the dichotomized group of mothers and PD were certainly related (*r* = 0.69), we previously ruled out their potential collinearity before its inclusion in the model as a covariate [See Supplementary *Methods* 4, Table S2]. All the analyses were performed with R (R Core Team, 2019).

*Emotional Availability (EA)* was measured in the context of mother-child free play using the EA Scale: Infancy to Early Childhood Version (Easterbrooks & Biringen, 2005). This scale operationalizes parental and child behavior in six scales that were factorized into one factor (EA) using Principal Component Analysis, given the existence of high inter-correlations among the scales. Two external observers blind to the mothers’ grouping made the ratings from the videos, and the inter-rater reliability of the ratings in each scale was adequate [see Supplementary *Methods* 5, Table S3 for the scales and testing]. EA was lower in neglectful (*M* = −0.62, *SD* = 0.92) than in control dyads (*M* = 0.71, *SD* = 0.48), *t*(43) = −6.18; *p* = .000; *δ* = 1.78, and it was used as a dependent variable.

*Alexithymia Scale*. Mothers completed the Toronto Alexithymia Scale (TAS-20) (Bressi et al., 1996; Taylor et al., 2003), which measures difficulty in identifying and verbalizing emotions, and minimizing of emotional experience by focusing attention externally. The 20 items are scored on a 5-point scale from strongly disagree to strongly agree. The TAS-20 has a 3-factor structure. Alexithymia was higher in NM than in CM in Factor Alex_DIF (*M* = 2.98; *SD* = 1.31; *M* = 2.18; *SD* = 1.07; *t*(43) = 2.21; *p* = .032; *δ* = .66), which assesses difficulty in identifying feelings; in Factor Alex_DVF (*M* = 3.35; *SD* = 1.3; *M* = 2.59; *SD* = 1.18; *t*(43) = 2.03; *p* = .048; *δ* = .60), which assesses difficulty in verbalizing feelings; and in Factor Alex_EOT (*M* = 3.05; *SD* = .61; *M* = 2.59; *SD* = .56; *t*(43) = 2.63; *p* = .011; *δ* = .78), which assesses externally oriented thinking. The overall mean difference was also significant (*M* = 3.10; *SD* = 0.88; *M* = 2.45; *SD* = 0.79; *t*(43) = 2.56; *p* = .013; *δ* = .77).

### Brain imaging

#### MRI acquisition

High-resolution T1-weighted MPRAGE anatomical volumes were acquired on a General Electric 3T scanner located at the University Hospital’s Magnetic Resonance Service for Biomedical Research at the University of La Laguna. A total of 196 contiguous 1mm sagittal slices were acquired with the following parameters: repetition time = 8.716 ms, echo time = 1.736 ms, field of view = 256 × 256 mm^2^, in-plane resolution = 1 mm × 1 mm, flip angle = 12.

#### Surface-based morphometry

CT and SA measures were obtained using FreeSurfer (version 5.1) (http://surfer.nmr.mgh.harvard.edu/). Surface cortical reconstruction included: motion correction, skull tripped, automated transformation into Talairach space, subcortical white matter (WM) and deep GM volumetric structure segmentation, triangular tessellation of the white surface (corresponding to the grey/white matter boundary) and pial surface (corresponding to the pia mater), and automated topology correction and surface deformation that optimally place the (inner) white surface and the (outer) pial surface (Dale, 1999; Fischl & Dale, 2000). Some deformations included surface inflation and a high-dimensional nonlinear registration to a spherical atlas. The segmentation and deformation algorithms produce representations of the CT that is the average of the closest distance from the white to the pial surface and from the pial to the white surface at each vertex on the tessellated surfaces. The SA is computed at the white surface, which is less sensitive to cortical thickness variations, and measured at each vertex as one-third of the area of each triangle that meets the vertex; in other words, it is the sum of the area of all the triangles that meet the vertex divided by three (Winkler et al., 2012). The CT and SA maps were smoothed using a 15 mm FWHM Gaussian filter.

### Statistical analysis for brain measures

Vertex-based analysis of covariance (ANCOVA) was performed to explore regional differences in CT and SA between CM and NM, adjusted for age. The model also included one regressor, described above as Psychiatric Disorders, to regress out the potential comorbidity with the negligence condition. Total intracranial volume (TIV) was also included as a nuisance covariate in the analysis of the SA. Monte Carlo simulation (10000 iterations) was used to perform cluster-wise correction of multiple comparisons, with an initial cluster-formation threshold of *t* = 2 (*p* < 0.01). Clusters were considered as significant at *p*-value < 0.05 corrected using Bonferroni. Regional differences were described based on the FreeSurfer Desikan/Killiany parcellation atlas (Desikan et al., 2006).

#### MNI to FreeSurfer surface coordinate projection

To make the cortical surface metric (CT/SA) results comparable with the GM volume results from the previous VBM study (Rodrigo et al., 2019), the *p*-value maps of the GM volumetric differences (CM > NM and NM > CM) were projected into the FreeSurfer average space (fsaverage). For the MNI to FreeSurfer surface coordinate systems projection, we used (Wu, 2018) standalone codes downloaded from the Github repository (https://github.com/ThomasYeoLab/CBIG/tree/master/stable_projects/registration/Wu2017_RegistrationFusion).

### Statistical analyses of brain structure-behavior associations

We followed two steps to determine the relationships between the cortical alterations observed in the vertex-wise analysis and all the behavioral variables. First, the CT and SA mean values for each mother, calculated from the clusters of difference between groups, were correlated with their EA scores in the play task, as well as with the Alexithymia scores (the overall and the three-factor scores) and Cognitive Integrity (CI). Next, mediation analyses were planned to test the potential mediation role of Alexithymia (the overall and the three-factor scores) and CI, between the cortical variables (CT/SA) and EA, but only in those cases in which significant correlations with both cortical and EA measures were obtained. Only two of the five behavioral variables fulfilled these conditions and were tested as mediators. For each mediation model the brain measures acted as the independent variable, Alexithymia/CI acted as a mediator – for model I and II, respectively – and EA acted as the dependent variable. Finally, as the majority of NM (16 out of 24) had suffered childhood maltreatment (neglect or physical abuse), they are more likely to be less sensitive when interacting with their child (Mielke et al., 2016). Therefore, we entered the maltreatment condition as a mediator to examine its potential contribution to the cortical - EA associations.

## RESULTS

### Altered CT and SA in neglectful mothers

Results in CT showed two distributed clusters with a pattern of thinner cortical grey matter in NM than in CM (top of Table 2). Each of these clusters spanned several contiguous anatomical regions in the right hemisphere (Figure 1). Cluster 1 involved a broad region of thinner right rostral middle frontal gyrus (herein RMFG), as part of the dorsolateral frontal gyrus (DLFG), with extension to the lateral and middle orbitofrontal area (OFC). Cluster 2 included a thinner right caudal anterior cingulate cortex (ACC) and a large midline region comprising rostral ACC and anterior /posterior middle cingulate cortex (MCC).

**Table 2.** Cortical thickness and Surface area differences, showing a thinner rostral frontal and orbitofrontal cortical cluster, a thinner anterior and posterior cingulate cortical cluster for neglectful mothers (NM < CM), a greater occipital lingual and fusiform surface cluster, and a greater caudal frontal, pars opercularis and precentral areas for neglectful mothers (NM > CM).

| Cluster/Regions | Total<br>Vertex | Cluster<br>*p*-value | Max<br>x, y, z (mm) | Max<br>-log<sub>10</sub>(*p*-<br>value) |
| --- | --- | --- | --- | --- |
| **Neglectful < Control Mothers (Cortical thickness)** |  |  |  |  |
| R. Rostral middle frontal,<br>Lateral and medial orbito-<br>frontal | 1880 | 0.014 | 21.1 51.5 -<br>12.5 | 3.44 |
| R. Caudal and rostral<br>anterior cingulate. Posterior<br>cingulate | 1838 | 0.039 | 5.7 18.9 26.9 | 3.04 |
| **Neglectful > Control Mothers (Surface area)** |  |  |  |  |
| R. Lingual, pericalcarine,<br>lateral occipital, fusiform | 3790 | 0.0002 | 7.4 -88.3 -11.3 | 3.35 |
| R. Caudal middle frontal,<br>pars opercularis, precentral | 2100 | 0.025 | 39.1 21 36.4 | 2.61 |

**Fig. 1.**
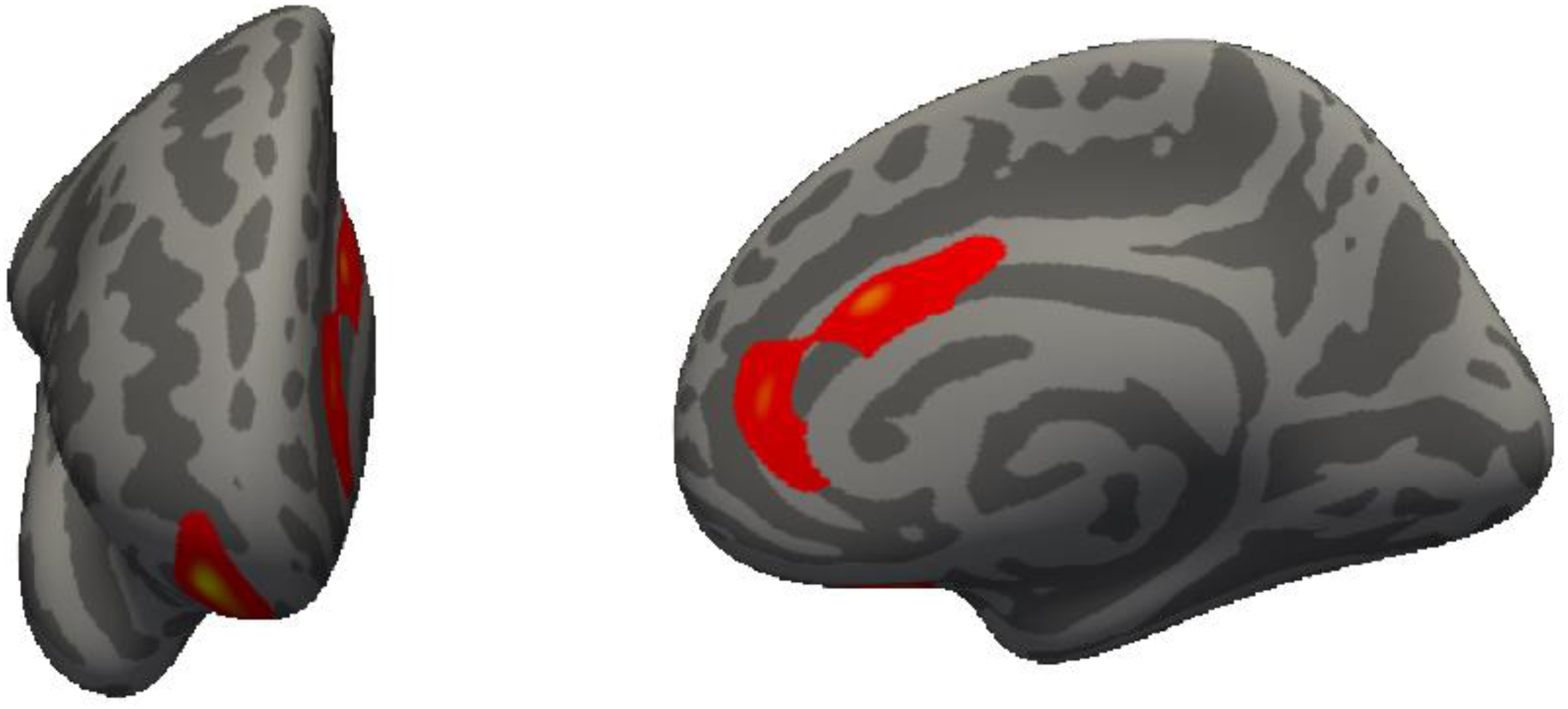
Frontal and cingulate regions on the right hemispheres showing decreased cortical thickness (CT) for neglectful mothers (NM) as compared to control mothers (CM). Background brain images represent the anterior (left image) and medial (right image) views of the right hemisphere inflated template. P < 0.05 corrected, age and Psychiatric Disorders measures were used as covariates, 10000 Monte Carlo interactions.

Regarding the SA, results showed two distributed clusters with a pattern of increased cortical SA in NM as compared to CM (bottom of Table 2, Figure 2). Cluster 1 involves a greater surface of the occipital right lingual, pericalcarine, lateral occipital and fusiform areas. Cluster 2 comprises a greater surface of the right caudal middle frontal gyrus (herein CMFG), the pars opercularis and the precentral areas.

**Fig. 2.**
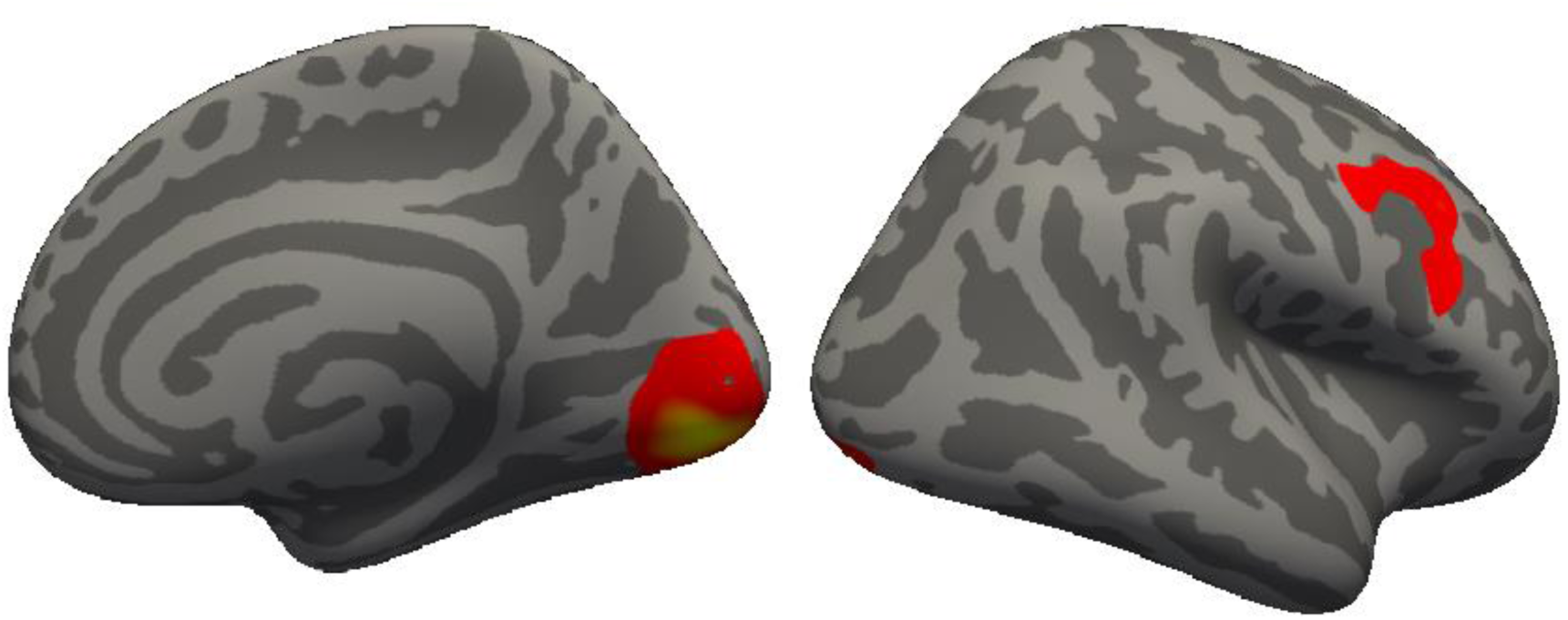
Occipito-temporal and frontal regions on the right hemispheres showing increased cortical surface area (SA) for neglectful mothers (NM) as compared to control mothers (CM). Background brain images represent the medial (left image) and lateral (right image) views of the right hemisphere inflated template. P < 0.05 corrected, age, total intracranial volume and Psychiatric Disorders were used as covariates, 10000 Monte Carlo interactions.

Comparisons between the current morphometric results and those obtained in our previous VBM study with a similar sample indicate that: (a) there is an overlap between CT and GM volumetric results showing reduced cortical thickness and volume inside the MCC for NM as compared to CM; and (b) SA and GM volumetric results coincide in reporting increased cortical SA and volume inside the fusiform and lingual areas for NM as compared to CM, although they did not overlap (Figure 3).

**Fig. 3.**
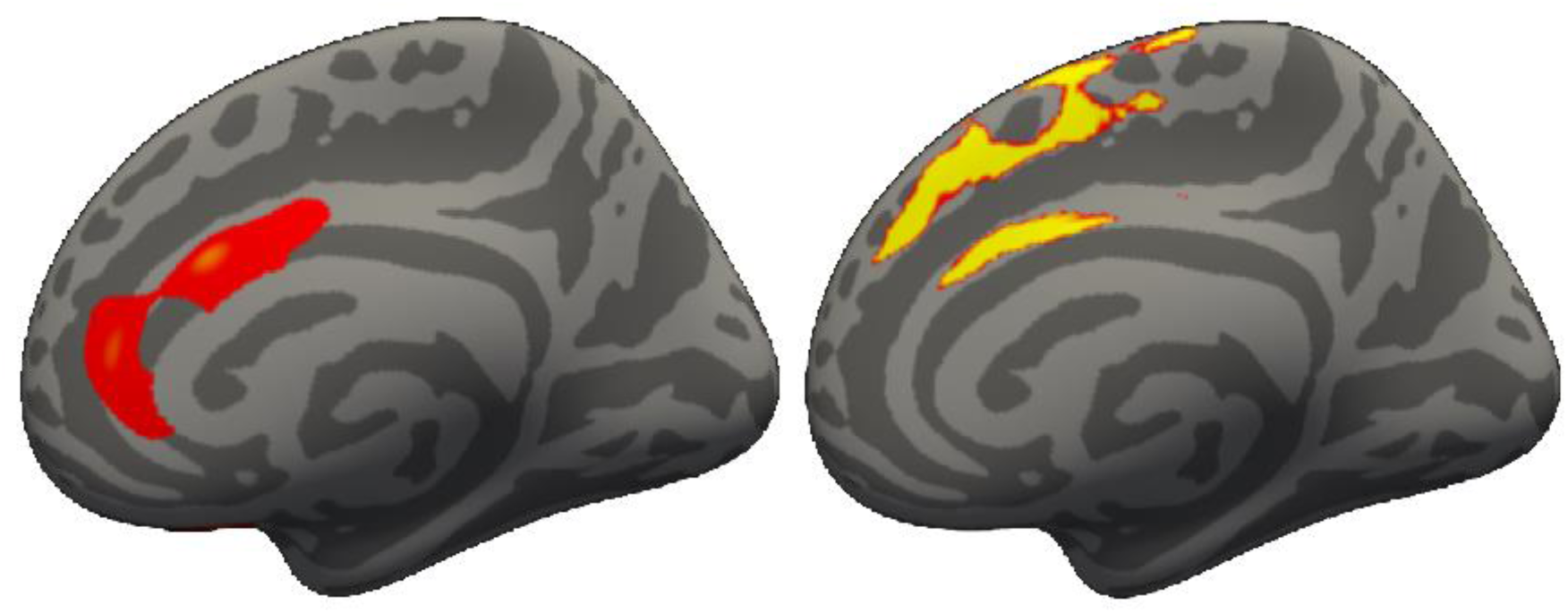
Coincidental decreases in CT and VBM found in the right middle cingulate cortex in neglectful mothers (NM) as compared to control mothers (CM). Background brain images represent the medial views of the right hemisphere inflated template for CT (left image) and VBM (right image). P < 0.05 corrected, age and Psychiatric Disorders measures were used as covariates, 10000 Monte Carlo interactions.

### Alexithymia and Cognitive Integrity as mediators of the effects of CT and SA on EA

No significant correlations were obtained between the EA and the mean values in the clusters of difference for either CT or SA. However, the psychological variables Alex_EOT and CI showed correlation in the right RMFG with the CT (for Alex_EOT: *r* = .33*; for CI: *r* = .40**) and with the EA (for Alex_EOT: *r* = -.30*; for CI: *r* = .61***), so they were suitable for the mediation analysis. Then, two mediation models were performed to test the role of Alex_EOT and CI as mediators in the relationship between the CT and EA (Figure 4). Model I (below) showed a significant direct positive effect of RMFG thickness on EA scores (average direct effect, *ADE* = 1.77, *p* = .004) and a negative effect through the mediation of the Alex-EOT (average causal mediation effects, *ACME* = −0.72, *p* = .036), indicating that the mediation of the External Oriented Thinking factor makes the relationship between the CT in the RMFG and the EA scores negative. Model II (above) showed a significant effect of RMFG thickness on EA through the positive mediation of CI (average causal mediation effects, *ACME* = 1.279, *p* = .01), indicating that CI mediates positively the relationship between RMFG thickness and EA scores. Using a bootstrap resampling procedure, in both models the parameters fell outside the confidence intervals, indicating that the results are not likely to be random. None of the meditation analyses trying to determine the possible role of the maltreatment condition in the relation between brain measures and EA showed significant results.

**Fig. 4.**
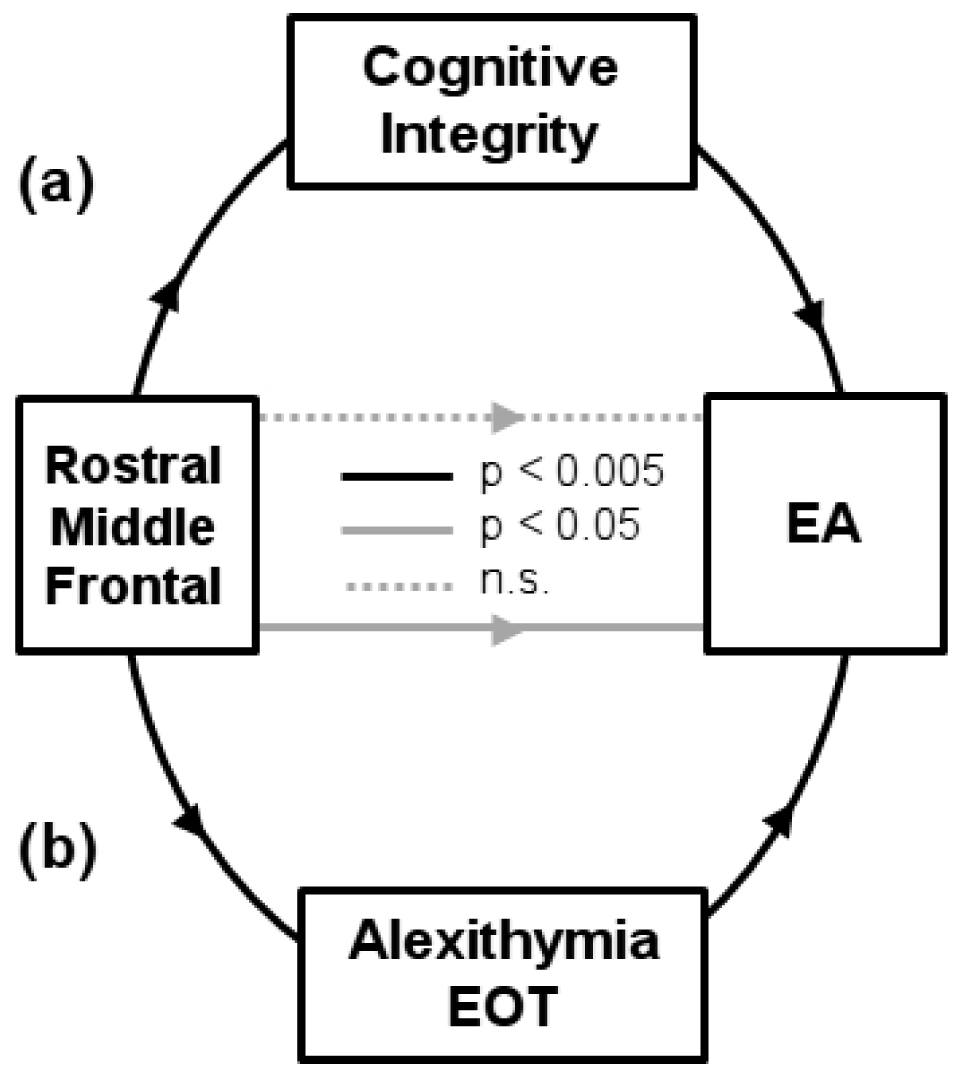
Mediation models in the CT of the Rostral Middle Frontal Gyrus on EA. Above (a) the model shows a positive relationship between Rostral Middle Frontal Gyrus and EA with no direct effects, only with the mediation of Cognitive Integrity. Below (b) the model shows a positive direct relationship of Rostral Middle Frontal Gyrus with EA that becomes negative if mediated by the Alexithymia factor of Externally Oriented Thinking.

## DISCUSSION

This study investigated, first, differential brain morphometric patterns in NM and CM and, second, their potential association with the mother-child bonding interaction (EA), adding evidence for brain structure-maternal caregiving associations. We found cortical alterations in NM using, for the first time in the study of this population, surface-based analysis. We also found that these cortical alterations were associated with EA via two psychological variables that influence the maternal capacity to effectively communicate emotions and to coordinate actions towards achieving joint goals during mother-child exchanges.

NM as compared to CM showed reduced CT in the right rostral middle frontal gyrus and orbitofrontal cortex (RMFG/OFC) and caudal anterior and middle cingulate regions (ACC/MCC) involved in the frontal-limbic modulation of cognitive control and in affective processing (Botvinick, 2007; Milham et al., 2001; Rudebeck & Murray, 2014). The anterior cingulate cortex plays a role in emotion regulation and social cognition (Etkin et al., 2011). NM also showed increased cortical SA in a region comprising occipital and fusiform areas specifically related to face processing (Fairhall & Ishai, 2008) and in the right caudal middle frontal gyrus. In previous studies, frontal and fusiform areas were found to undergo critical morphometric adaptations (CT increases) in the postpartum period in a normal range of sensitive mothering (Kim et al., 2018) and to suffer volumetric alterations (GM decreases) in neglectful mothering (Rodrigo et al., 2019), highlighting the relevance of these areas to support variations in the quality of maternal caregiving.

The CT and SA alterations in NM were compatible with the GM volumetric patterns obtained in a previous VBM study (Rodrigo et al., 2019), providing evidence of the correspondence between GM volume, CT and SA measures (Hogstrom et al., 2012; Winkler et al., 2010). Specifically, the cortical thinning in the MCC overlapped with the GM volume reduction obtained in the same area, whereas the cortical thinning in the middle frontal gyrus area can be associated with the GM volume reduction obtained in an adjacent frontal region (i.e., the inferior frontal gyrus). In turn, the cortical SA increase in occipito-temporal areas was in parallel to, though it did not overlap with, the GM volume increase in the right fusiform and the cerebellum.

Interestingly, this reverse pattern of GM increase in occipito-temporal regions and decrease in frontal regions is replicated through the different measures. A similar reverse pattern was obtained in the oscillatory rhythms of NM in response to emotional stimuli when compared to CM (León et al., 2014): the higher the increase in theta band oscillations (and lower alpha) at occipital sites, the lower it was in the frontal sites. Both morphometric and oscillatory patterns seem to reflect a lower engagement of frontal regulatory areas – relevant in sensitive caregiving – over the occipital areas in NM. The lower fMRI pattern of activation in response to crying faces in occipital areas in NM as compared to CM (León et al., 2019) may also reflect a lower top-down regulation from frontal regions. Further studies are needed to determine the intervening factors that explain the intricate interactions between the dynamic and structural (more static) alterations in regions involved in sensitive caregiving.

Notice that the cortical thinning in the right RMFG and OFC can be related to the susceptibility of these areas to neurodegeneration (Seeley et al., 2009) specifically related to age (Buckner et al., 2000). However, the influence of age can be ruled out here, since the analysis was controlled for age effects and the neglectful mothers were slightly younger than the control mothers. Besides, substantial macrostructural WM volume reductions in bilateral frontal areas in NM (Rodrigo et al., 2019) accompanied the frontal and cingulate cortical thinning observed in NM in this study. This convergence is congruent with studies reporting that alterations in white matter can influence CT values in the altered white matter regions (Belathur Suresh et al., 2018). Taken together, our results suggest that the cortical thinning observed in NM could be a result of accelerated/truncated pruning (Goldstone et al., 2018) and/or a concurrent loss of myelinated white matter that may have occurred during development (Mills & Tamnes, 2014). Nevertheless, the direction of the relationship between CT and myelination is not yet clear, solely based on the morphometric measures.

In the analysis of the brain structure-behavior associations, we found a mediated association, via alexithymia and cognitive integrity, between the rostral middle frontal gyrus (RMFG) and the EA exhibited in a mother-child interactive play task. Several models suggest that the RMFG involves hierarchical and non-stimulus-dependent levels of functioning, which is typical of high regulatory instances (Badre & D’esposito, 2009; Bahlmann et al., 2014; Botvinick, 2008). Accordingly, we did not find evidence of a direct RMFG-EA association but of an association mediated by both emotional and cognitive psychological traits: alexithymia and cognitive integrity. Previous studies have shown that frontal areas are also related to individual differences in trait alexithymia (Lenzi et al., 2013) and cognitive integrity (Arlt et al., 2013).

In our study, higher scores in the alexithymia factor of externally oriented thinking, consisting of a tendency to focus on the simpler and external aspects of the events rather than their psychological correlates (Taylor & Bagby, 2004), negatively mediated the association of RMFG to EA. This external focus of trying to avoid the complexity of the affective dimension may entail difficulties for the appropriate self-awareness and regulation of the mother’s own emotions in the course of communicative dyadic exchanges. Also, a positive RMFG association with EA was mediated by higher efficiency in cognitive processing, which may facilitate more organized and sustained courses of action during ongoing exchanges. This may increase, in turn, the child’s ability to predict the mother’s actions, leading to a better mother-child synchronic functioning (Beebe & Steele, 2013). Previous evidence has shown that mothers with a lower working memory capacity exhibited poorer cognitive control over their thoughts and behaviors during a cooperation task with their child (Deater-Deckard et al., 2010). As a result, to the extent that NM show alterations in the RMFG, enhanced by a tendency toward higher alexithymia and poorer cognitive functioning, their ability to sensitively respond to and effectively organize their behavior towards the child may be affected. These findings support the view that executive functions are required for the mothers’ own emotion, cognitive and behavior regulation towards the child (H. J. Rutherford et al., 2015).

The results of this study should be considered in light of two limitations. First, there is some overlap in the present sample between being neglectful, exhibiting greater psychiatric vulnerability and having suffered childhood maltreatment, as corresponds to the typical risk profile in this population. Building on the neural differences found in this study, future research with larger samples would allow for an orthogonal design crossing NM and CM with psychopathological conditions and own childhood maltreatment to determine the respective contribution of these different factors to the neural alterations associated with maternal neglect. Second, the cross-sectional design used does not allow us to infer whether the neurological alterations have causal relationships with EA.

In conclusion, this study has revealed specific cortical feature alterations, mainly in frontal, cingulate and fusiform cortices, in NM that can reflect the potential failures in high-level regulatory functioning that are relevant for mother-child interactive bonding. These cortical alterations, along with the poor viso-emotional processing demonstrated by ERP and oscillatory measures (León et al., 2014; Rodrigo et al., 2011), the reduced fMRI activation in frontal and cingulate regions for crying faces (León *et al*., 2019), and the poor white matter connectivity in the viso-emotional circuits (Rodrigo *et al*., 2016), provide a more complete picture of the neurological underpinnings associated with neglectful caregiving and their impact on mother-child bonding. In this characterization we see alterations emerge in regions related to both the automatic “intuitive parenting” responsible for the fast and empathic response to the infant’s emotional signals, as well as in other areas more related to the “regulated parenting” responsible for maintaining a self-regulated emotional state, a shared focus of attention and a temporal coordination and contingency with the child’s actions. Both modalities of parenting should be perfectly attuned to establish a positive mother-child interactive bonding. The outputs of this study can be instrumental in the development of imaging-based diagnostic classification and predictive models of maternal neglect. These achievements can help improve effective prevention and intervention strategies to decrease the risk of neglectful mothering and promote infant health and wellbeing.

## Acknowledgments

We thank the Health and Social Services staff and all the mothers and their children who participated in this study.

