## SUPPLEMENTARY MATERIAL for "Cortical surface alterations in cingulate and frontal regulatory areas underlie insensitive mothering"

**TITULO**
**SUPPLEMENTARY MATERIAL**

### Methods

#### 1) Participant recruitment strategy and procedure

All mothers in the neglectful group exhibited the three main subtypes of neglect and scored positively on all indicators: physical neglect (inadequate food, hygiene, clothing, and medical care), lack of supervision (child is left alone or in the care of an unreliable caregiver) and educational neglect (lack of cognitive and socioemotional stimulation and lack of attention to child's education). After the selection of the sample, social workers reported on the sociodemographic and neglect risk profile and asked mothers for permission to be contacted by phone. Those mothers who gave permission were contacted by our collaborator and were informed about the general goal of the study (to participate in a study about mother-child relationships avoiding the use of the term neglect in any case) and the procedure to be followed upon their acceptance. Then, the collaborator picked them up at their homes at their convenience to bring them to the scanning session at the Hospital where they gave their informed written consent and passed the MRI sequence under a resting state condition without stimuli being presented. In a second session carried out at their homes, the same collaborator collected the mothers' response to the questionnaire, gave a gift to the child and video recorded the mother-child play interaction. At the end of the session mothers received a monetary compensation.

#### 2) Risk profile measures

Social workers reported on a series of risk indicators (presence: 1; absence: 0) that are commonly used to assess maternal neglect. *History of abuse/neglect* refers to whether mothers have suffered childhood maltreatment (either abuse or neglect) in their own history (scoring 1); *Intimate partner conflict* refers to whether mothers are experiencing overt conflictive relationships with their partner (scoring 1); *Chronic physical illness* refers to whether they are currently experiencing poor health conditions permanently or very frequently (scoring 1); *Poor household management* refers to whether the home is dirty and/or untidy, with irregular meals and/or dirty clothing (two is enough for scoring 1); *Disregard health/education needs* refers to lack of or discontinuous medical checks, irregular vaccines, and/or poor support for learning (two is enough for scoring 1); *Disregard emotional/cognitive needs* refers to poor attention to the child's emotional expressions and/or lack of response to infant curiosity (one is enough for scoring 1); *Rigid/inconsistent parental norms* refers to an application of rules without taking into account the childrearing situations and/or arbitrary changes to norms applied to the same situations (one is enough for scoring 1).

**3) Table S1. Psychopathological conditions stratified by Group**

| | Neglectful group ($n$ = 25) *M* (*SD*) | Control group ($n$ = 23) *M* (*SD*) | *t*(46) | Effect size δ |
|---|---|---|---|---|
| *Major Depressive Episode* | 1.79 (2.4) | 0.23 (0.5) | 3.04** | 0.91 |
| Dysthymia | 1.5 (2.2) | 0.3 (0.5) | 2.54* | 0.75 |
| Suicidality | 0.4 (0.7) | 0 | 2.65* | 0.79 |
| *Hypo/Manic Episode* | 1.8 (2.1) | 0.1 (0.3) | 3.79** | 1.14 |
| *General Panic Disorder* | 6.4 (5.5) | 0.8 (2.2) | 4.57*** | 1.36 |
| Agoraphobia | 0.6 (8) | 0.3 (0.6) | 1.24 | 0.37 |
| Social Phobia | 0.5 (0.9) | 0 | 2.62* | 0.78 |
| Obsessive-Compulsive | 1.1 (1.5) | 0.2 (0.5) | 2.62* | 0.78 |
| Post-traumatic Stress Disorder | 1.6 (2.3) | 0.7 (0.7) | 1.61 | 0.48 |
| Alcohol Dependence/Abuse | 0.9 (0.2) | 0.4 (0.4) | 0.51 | 0.16 |
| Drug Dependence/Abuse | 0.12 (0.3) | 0 | 1.69 | 0.50 |
| Psychotic Disorders | 0.6 (1.2) | 0.2 (0.5) | 1.20 | 0.36 |
| Bulimia Nervosa | 0 | 0 | 1 | 0.32 |
| *Generalized Anxiety Disorder* | 3.2 (3.2) | 0.7 (0.9) | 3.53*** | 1.06 |
| *Antisocial Personality* | 1.2 (1.2) | 0.1 (0.3) | 3.8*** | 1.12 |

*$p$≤05; **$p$≤.01; ***$p$≤.001; Italic Fond in some variables indicates those that survived the Bonferroni test in the Group comparisons, and were submitted to a Principal Component Analysis to obtained the variable Psychiatric Disorders.

**4) Table S2**. **Collinearity indexes between the Group (as a dichotomic variable) and the psychopathological conditions, both as a factor PD and separately as individual disorders (within brackets are the cutoff values for non-collinearity).**

| | VIF (<10) | TOL (>0.30) | CN (<10) | Shared Variance (<0.50) |
|---|---|---|---|---|
| **Factor score "Psychiatric Disorder"** | **1.88** | **0.53** | **2.31** | **0.46** |
| Major Depressive Disorder | 1.19 | 0.83 | 1.53 | 0.16 |
| Hypo/Manic Episode | 1.29 | 0.77 | 1.67 | 0.22 |
| General Panic Disorder | 1.44 | 0.69 | 1.86 | 0.30 |
| Generalized | 1.25 | 0.79 | 1.63 | 0.20 |

| | | | | |
|---|---|---|---|---|
| Anxiety Disorder | | | | |
| Antisocial Personality | 1.29 | 0.77 | 1.68 | 0.22 |

Note: VIF: Variance Inflation Factor, TOL: Tolerance, CN: Condition Number.

We assessed the potential Multicollinearity (MCL) calculating three well-known indexes: the Variance Inflation Factor (VIF), the Tolerance (TOL), and the Condition Number (CN). We calculated them for the overall factor PD with the Group, as well as separately for each psychiatric disorder that survived the Bonferroni test in the group comparisons (Table S2). We also measured the shared variance of the psychiatric variables with the Group (last column). According to the literature, the general criterion for non-collinearity is a Variance Inflation Factor (VIF) less than 10, a Tolerance greater than 0.30, a Condition Number less than 10, and a shared variance less than 0.50 (Belsley et al., 2005; Kovács et al., 2005; Tabachnick & Fidell, 2007). Our results (see Table S2) showed that all the values fall below the corresponding cutoffs for collinearity. That made it possible to include the PD as a covariate together with Group in the SPM model, to control as much as possible its effect on the results.

**5) Table S3. Inter-rater reliabilities and one-factor component loadings of the Emotional Availability Scales.**

| | Kappa coefficients | Component loadings |
|---|---|---|
| Sensitivity | 0.94 | 0.892 |
| Structuring | 0.90 | 0.933 |
| Nonintrusiveness | 0.87 | 0.830 |
| Nonhostility | 0.92 | 0.767 |
| Responsiveness | 0.92 | 0.892 |
| Involvement | 0.86 | 0.867 |

For the emotional availability score, the mother-child interaction was videotaped at home, in the context of mother-child free play, at the moment when the family received a toy as a gift for participation in the study. Mothers were instructed to use the toy and play with the child as they usually do. Ratings from the videos were based on the Emotional Availability Scale, which operationalizes four aspects of parental behavior: *Sensitivity* (9 points) - the parent shows contingent responsiveness to child signals; *Structuring* (5 points) - the parent appropriately facilitates the child's play; *Non-intrusiveness* (5 points) - the parent is able to support the child's play without being over directive and/or interfering; *Non-hostility* (5 points) - the parent is able to behave with the child in a way that is not rejecting or antagonistic. The scale also measures two aspects of child behavior: *Responsiveness* (7 points) - the child's ability and interest in exploring on his or her own and in responding to the parent's bids; *Involvement* (9 points) - the child's ability and willingness to engage the parent in interaction (Table S3). To obtain a more simple structure of the six standardized scales, a Principal Component Analysis was performed. The result yielded a single factor structure: KMO = 0.84, Eigenvalue = 4.49, with an explained variance of 75%.
